# Predictability of Sleep Slow Oscillation Emergence and Spatial Extent from Pre-Onset Neural Dynamics

**DOI:** 10.64898/2026.01.09.698723

**Authors:** Mahmoud Alipour, Wim van Drongelen, Paola Malerba, Joel Voss, David Satzer

## Abstract

Slow oscillations (SOs; ∼0.5–1.5 Hz) are a hallmark of non-rapid eye movement (NREM) sleep and are known to support memory consolidation and large-scale cortical communication. Although their instantaneous dynamics are well characterized, the neural processes that precede SO initiation—and whether they predict the spatial extent of the upcoming event—remain largely unknown, despite their potential utility for anticipatory closed-loop intervention. Using high-density EEG from 29 healthy adults, we examined neural activity in the 2-s interval preceding the SO trough. Analyses focused on distinct SO subtypes defined by their spatiotemporal properties: Global, Frontal, and Local. We specifically focused on Global and Frontal events, both of which originate frontally but differ in their propagation. We quantified instantaneous spectral power, time–frequency dynamics, phase–amplitude coupling (PAC), and amplitude–amplitude coupling. Across all analyses, theta-band power (4–8 Hz) emerged as the earliest and most robust predictor of SO initiation, remaining informative after restricting analyses to temporally isolated SOs to reduce potential residual-aftereffect confounds from preceding SOs. Theta power exhibited a sustained rise beginning nearly two seconds before the SO trough (∼1.6 s before onset). Theta power increases were absent in surrogate epochs and reliably differentiated Global from Frontal SOs with moderate-to-strong effect sizes (Cohen’s d = 0.45–0.77), demonstrating the robustness of theta power as a physiological signature. Mechanistically, Global SOs were preceded by enhanced delta–theta PAC and broad low-frequency synchronization, whereas Frontal SOs were preceded by elevated theta/alpha-to-beta/low-gamma coupling, reflecting a state of locally enhanced coupling at frequencies higher than the SO range, which appears to restrict propagation. A simple logistic regression classifier using only pre-onset theta power achieved >95% accuracy in distinguishing SOs from surrogate events and differentiated Global from Frontal SOs with ∼83% multiclass accuracy, showing further sensitivity improvements when delta power was included. These findings demonstrate that isolated SOs are preceded by structured network dynamics that are tied to their spatial extent. Thus, theta activity can be used to predict SO occurrence and might be leveraged in closed-loop neuromodulation.

## 1. Introduction

Slow oscillations (SOs; ∼0.5–1.5 Hz) are the hallmark of non–rapid eye movement (NREM) sleep and reflect a fundamental bistability of neurons in the thalamocortical system during NREM, in which neuronal populations alternate between periods of quiescence (down-states) and synchronized firing (up-states)^1,2^. These large-amplitude, slow fluctuations orchestrate neuronal activity across distributed cortical networks and play a central role in memory consolidation, synaptic rescaling, and brain homeostasis^3–5^. Given their importance for sleep-dependent cognitive and physiological functions, SOs have become a primary target for closed-loop neuromodulation approaches aimed at enhancing cognition and treating neurological disorders^6,7^.

Importantly, SOs are not a homogeneous phenomenon. Although SOs predominantly originate in the prefrontal neocortex, their spatial propagation across the scalp varies considerably^8–10^. Based on their spatiotemporal propagation patterns, SOs can be classified into distinct subtypes, referred to as Global, Frontal, and Local events^11^. In this classification, Frontal SOs remain confined to anterior cortical areas, whereas Global SOs propagate as traveling waves across widespread cortical regions in an anteroposterior direction^11^. Other classification systems further characterize this heterogeneity using waveform morphology and synchronization efficiency, distinguishing “Type I” slow waves—which exhibit steeper, larger, and more widespread down-state transitions—from “Type II” waves, which are smaller, more localized, and possess flatter transitions^12,13^. These morphological distinctions likely overlap with the Global/Frontal dichotomy, suggesting the coexistence of multiple synchronization processes during NREM sleep. The distinction of SOs based on their spatiotemporal propagation has functional implications. Previous research demonstrated that among the different subtypes, Global SOs are strongly associated with episodic memory, whereas spatially restricted SOs show little to no such association^14^. Furthermore, evidence indicates that Global SOs preferentially coordinate with sleep spindles—a coupling mechanism strongly linked to memory consolidation—while non-Global SOs show significantly weaker associations with this process^11^. The spatiotemporal organization of SOs may also serve as a biomarker for health and disease. For instance, a recent study by our team demonstrated that the proportion of SO subtypes shifts in pathology; specifically, individuals with hypersomnolence exhibit a significantly higher percentage of Frontal SOs compared to healthy controls^15^. Furthermore, in other pathological conditions, such as epilepsy, SOs can be co-opted by interictal epileptiform activity^16–18^. Consistent with the idea that spatial expression matters, intracranial recordings indicate that interictal spikes occur preferentially during high-amplitude, widespread slow waves. This suggests that SO extent may index a specific vulnerability to epileptiform recruitment^16^. This phenomenon raises a critical concern: indiscriminate enhancement of SOs may inadvertently amplify pathological dynamics rather than beneficial physiological rhythms. Consequently, effective therapeutic modulation requires more than simple event detection; it demands the precise discrimination of SO subtypes and their spatial extent to ensure safe and targeted intervention.

A critical limitation of existing closed-loop stimulation systems that aim to enhance SOs during NREM sleep is their reliance on ‘reactive detection’ of SOs^19^. Most algorithms identify SOs only after the oscillation has begun—for example, upon crossing a negative voltage threshold—introducing delays that are further compounded by signal filtering and hardware latencies^6,19^. Because the SO up-state, typically the target of excitatory stimulation, lasts only a few hundred milliseconds, these delays often lead to suboptimal or mistimed intervention^6^. Moreover, current approaches focus analytical attention on the oscillation itself rather than on the neural state that precedes it. This severely impedes any closed-loop approach from predicting SO events early enough to target them with accuracy. Of note, recently Menicucci et al. (2015) demonstrated that fluctuations in sigma activity (defined as 12–18 Hz in their study) precede SO onset and correlate with the amplitude and slope of the upcoming wave, suggesting the presence of a distinct thalamic preparatory phase^9^. Converging evidence from intracranial and scalp recordings reported by Gonzalez et al. (2018) further suggests that theta-band activity (defined as 5–8 Hz in their study) can emerge prior to cortical and thalamic down-states and may contribute to SO initiation^20^. More broadly, large-scale studies of human sleep EEG demonstrate that slow-wave properties depend on the pre-existing network state, consistent with the presence of preparatory dynamics preceding SO events^12^. Together, these findings suggest that activity preceding SO events may be leveraged as predictors of SO emergence. If SOs arise from such identifiable preparatory dynamics, then detecting precursors could enable a shift from ‘reactive’ to ‘predictive’ stimulation strategies, allowing stimulation to be optimally timed to forthcoming SOs while bypassing detection delays altogether.

In this study, we tested the hypothesis that SOs are preceded by structured neural dynamics that predict not only their emergence but also their subsequent spatial extent. Using high-density EEG recordings from 29 healthy adults in the ANPHY-Sleep database^21^, we applied an established detection and clustering framework to identify SOs and classify them into Global and Frontal subtypes based on their spatiotemporal co-occurrence patterns^11^. We then shifted the analytical focus from the SO itself to the two-second window preceding the SO trough, quantifying a comprehensive set of pre-onset features including instantaneous spectral power, phase–amplitude coupling, and amplitude–amplitude coupling. Analyses were centered on prefrontal and frontal regions, which show the earliest scalp manifestations of SOs and from which both Frontal and Global events diverge.

Our findings demonstrate that SOs emerge from distinct preparatory brain states. We identify a robust increase in theta-band activity beginning nearly two seconds before SO onset, which reliably predicts upcoming SOs with high accuracy when compared to surrogate events. Critically, this pre-onset theta enhancement differentiates Global from Frontal SOs, indicating that spatial propagation is informed by activity preceding the unfolding of the SO event. Beyond power alone, we show that Global SOs arise from a neural state characterized by enhanced low-frequency coordination, including strengthened delta–theta coupling, whereas Frontal SOs emerge from a more locally synchronized state with prominent coupling at higher frequencies. Together, these results establish a physiological basis for predictive closed-loop neuromodulation and identify theta-band dynamics as a principled biomarker for selectively targeting widespread, functionally relevant SOs.

## 2. Materials and Methods

### 2.1. Dataset

We analyzed full-night high-density EEG recordings from the open-access ANPHY-Sleep database, which includes 29 healthy adults (13 female; age 32.17 ± 6.30 years)^21^. EEG was acquired using an 83-channel 10–10 montage at 1000 Hz, accompanied by EOG and EMG channels for manual sleep staging based on the American Academy of Sleep Medicine criteria^22^. Sleep scoring distinguished Wake, N1, N2, N3, and REM sleep 30-second epochs. Mastoid channels (M1/M2) were used during sleep staging but were not included in the publicly released dataset. A full description of data collection and scoring procedures is provided in the original database publication^21^.

### 2.2. Preprocessing

All preprocessing and analyses were performed in MATLAB (R2025b, MathWorks, Natick, MA, USA) using custom scripts developed by the authors. EEG data were downsampled to 250 Hz. The supraorbital (SO1/SO2) and zygomatic (ZY1/ZY2) electrodes, although categorized as EEG channels in the released dataset, were removed from our analysis to minimize ocular and facial EMG contamination. Because standard mastoid references (M1/M2) were not available, we implemented a contralateral mastoid–proxy referencing scheme using TP9 and TP10, the temporal electrodes closest to the mastoids. This approach preserves left–right symmetry and has been validated in prior studies as an effective proxy for standard mastoid referencing^23,24^.

Artifact removal followed a multi-stage procedure combining automated detection with statistical criteria. The dataset included an artifact matrix identifying gross artifactual 30-s epochs across electrodes, generated using the High-Density-SleepCleaner algorithm—an open-source, semi-automatic, GUI-based routine that detects channel- and epoch-specific artifacts using multiple signal-quality markers, including delta and beta power and broadband amplitude outliers^21,25^. Residual muscle-related contamination was addressed using the method described by Brunner and colleagues, which flags bins in the 26.25–32 Hz band whose amplitude exceeds four times the median of surrounding bins^26^, as well as the broadband (4–50 Hz) procedure of Wang and colleagues, which excludes 5-s bins exceeding six times the global median^27^. In addition, epochs with kurtosis values greater than 5 or amplitude deviations exceeding 5 standard deviations were removed. Prior to spectral analyses, a 60-Hz notch filter (2-Hz bandwidth, second-order Butterworth, bidirectional) was applied. All filtering and feature extraction steps were performed offline using non-causal processing to enable precise characterization of pre-onset neural dynamics; causal real-time implementations were not evaluated in this study.

### 2.3. Slow Oscillation Detection and Spatiotemporal Categorization

SOs were identified during stages N2 and N3 using established criteria^10,11,28^, adjusted to account for the reduced absolute amplitudes introduced by the TP9/TP10 proxy mastoid reference. EEG signals were band-pass filtered between 0.1–4 Hz using a zero-phase, second-order Butterworth filter, and SO candidates were defined as half-waves between consecutive positive-to-negative and negative-to-positive zero crossings.

To qualify as SOs, candidate events were required to exhibit both a sufficiently deep negative deflection and adequate overall excursion. Specifically, the negative peak had to reach at least −55 µV, and the peak-to-peak amplitude had to be no smaller than 80 µV. Given the attenuated amplitude resulting from TP9/TP10 referencing, negative peak threshold was set lower than those typically reported using true mastoid references^15,29,30^. Temporal criteria further ensured physiological plausibility, requiring the negative half-wave to last between 0.25 and 1 s and the full cycle between 0.5 and 5 s. These parameter ranges were selected after sweeping multiple threshold combinations and retaining those that preserved canonical physiological gradients—such as decreasing SO density and amplitude along the anteroposterior axis—and propagation dynamics^10,15,29^.

To characterize the spatial extent of SOs, each detected event was classified as Global, Frontal, or Local based on its co-occurrence pattern across EEG channels using the established algorithm introduced by Malerba et al. (2019)^29^. This approach relies on unsupervised clustering of binary channel co-occurrence patterns. Following this prior work, we set the number of clusters to k = 3, consistent with the established implementation of this algorithm and corresponding to the point at which further increases in k do not yield qualitatively distinct or more stable spatial patterns. For each SO, a binary co-occurrence matrix was constructed by identifying electrodes whose troughs occurred within a 400-ms window of one another. K-means clustering using Hamming distance (k = 3, 200 replicates) was applied to these binary co-occurrence vectors. Cluster centroids were mapped to scalp topographies and used to assign subtype labels. Global SOs exhibited widespread bilateral engagement across frontal, central, and posterior regions; Frontal SOs were largely restricted to prefrontal and frontal electrodes; and Local SOs occurred sparsely without a coherent spatial pattern.

Only isolated SOs were included in subsequent analyses. Events occurring within 1.5 s of a preceding SO trough were excluded to avoid contamination from SO trains, which could otherwise distort or confound pre-onset spectral dynamics. This exclusion was applied after SO subtype classification to preserve consistency with the original implementation of the clustering algorithm and prior work. The total number of detected SOs, subtype-specific SO counts, and the number of SOs excluded due to train removal (both overall and per subtype) are reported in Supplementary Table S1. This train-exclusion criterion resulted in the retention of 73% of detected SO events.

### 2.4. Surrogate Epoch Construction

To dissociate SO-specific dynamics from background NREM activity, we generated surrogate control datasets for each subject. Surrogate epochs were created by randomly sampling time points from N2 and N3 sleep, ensuring that each epoch occurred at least ±3 s away from any detected SO trough to avoid contamination. Crucially, the number of surrogate epochs was matched to the number of identified Global and Frontal SOs within each subject. This ensured that the signal-to-noise ratio of the surrogate average was comparable to that of the real SO conditions. Surrogate events underwent identical processing steps as real SOs—including power extraction, smoothing, and baseline normalization—yielding a robust null distribution for comparison.

### 2.5. Quantification of Neural Dynamics Preceding SO events

The following sections describe the signal processing pipelines used to quantify pre-onset neural activity. We adopted a hierarchical analysis framework, first quantifying instantaneous spectral power to identify specific frequency bands showing significant pre-onset activation. Based on these identified active bands, we then applied cross-frequency coupling to characterize the network interactions—such as low-frequency coordination—that could organize the pre-onset state. Finally, time–frequency decomposition provided a high-resolution validation of these spectral dynamics, while surrogate control epochs were generated to establish a rigorous baseline for statistical comparison.

#### 2.5.1. Instantaneous Power Extraction

To characterize pre-onset power dynamics, raw EEG was filtered into frequency bands (delta 0.5–4 Hz, theta 4–8 Hz, alpha 8–12 Hz, sigma 12–16 Hz, beta 16–30 Hz, low gamma 30–59 Hz, high gamma 61–119 Hz) using zero-phase, second-order Butterworth filters. Instantaneous power was obtained by computing the analytic signal via the Hilbert transform and squaring its magnitude to yield the envelope |z(t)|², providing a continuous, phase-independent estimate of oscillatory energy^31^. To emphasize sustained trends rather than momentary fluctuations, the instantaneous power time series was smoothed with a 500ms sliding window advancing in 50ms steps. Power values were expressed as percent change relative to a baseline defined as 2-3 s before the SO trough.

#### 2.5.2. Cross-Frequency Coupling

Phase–amplitude coupling (PAC) quantified the influence of low-frequency phases on higher-frequency amplitude fluctuations^32^. For each SO and surrogate epoch, phases were extracted using Hilbert transforms of band-pass–filtered signals (delta through low gamma), and amplitude envelopes were extracted from higher-frequency bands (theta through high gamma). The Modulation Index (MI) provided a normalized measure of coupling strength and was computed as an adapted Kullback–Leibler (KL) divergence of the phase–amplitude distribution from uniformity^33^. MI was calculated by first binning the phase of the low-frequency oscillation into N = 18 (each 20^°^) equally spaced bins and determining the mean amplitude of the high-frequency envelope within each phase bin to construct a discrete phase–amplitude distribution, P. This distribution was then compared with a uniform distribution U = 1/N using the KL divergence, defined as:

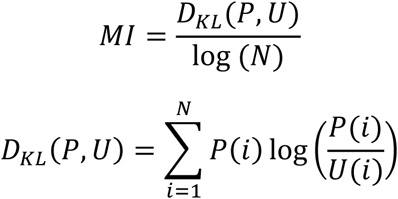

This normalization bounds MI between 0 (no coupling) and 1 (extreme coupling)^33^. The resulting MI values were additionally baseline-normalized by subtracting the MI computed in the matched non-event baseline window. Amplitude–amplitude coupling (AAC) was computed as the Pearson correlation coefficient between the Hilbert amplitude envelopes of all frequency band pairs, capturing co-fluctuations in broadband activity^34^.

#### 2.5.3. Time–Frequency Decomposition

Time–frequency representations were computed in FieldTrip (version 20251201) using time-frequency analysis functions with Hanning tapers^35^. A sliding window with a frequency-dependent length of 4 cycles provided 0.5 Hz spectral resolution from 1 to 119 Hz. Analyses spanned −4 to +3 s relative to the trough, with baseline 2-3 s before SO trough and statistical comparisons restricted to −2 to +2 s relative to the trough to avoid edge artifacts.

### 2.6. Statistical Analysis

Given that SOs predominantly originate in and propagate through prefrontal and frontal regions^8^, this study focused on Global and Frontal subtypes to specifically characterize the dynamics distinguishing widespread propagation from frontally confined activity. Local SOs were excluded from the predictive modeling analyses due to their lack of coherent spatial propagation patterns. To represent the primary hub of SO generation while maximizing signal-to-noise ratio, we constructed a “Broad Frontal” virtual channel. All spectral and coupling analyses were performed on this composite signal, calculated as the average of the signals from 17 prefrontal and frontal electrodes (Fp1, Fp2, Fpz, AF3, AF4, AF7, AF8, AFz, F1–F8, Fz). This spatial averaging approach reduces the influence of channel-specific artifacts and provides a robust estimate of frontal cortical dynamics relevant to SO initiation.

Statistical comparisons followed a metric-specific protocol. For spectral power, expressed as percent change from baseline (−3 to −2 s), deviations from baseline were assessed using one-sample t-tests against zero. Cross-frequency coupling measures (PAC and AAC) were compared to their respective baseline windows using paired t-tests. Direct within-subject comparisons between SO subtypes (Global vs. Frontal) and between real and surrogate events were also performed using paired t-tests. Because the number of surrogate trials was matched to the number of real events per subject, this approach rigorously compared event-locked dynamics against the representative background activity level of the same subject.

Multiple comparisons for PAC and AAC matrices were controlled using the Benjamini–Hochberg false discovery rate (FDR; α = 0.05)^36^. Time-frequency contrasts were evaluated in FieldTrip using cluster-based permutation testing (Monte Carlo method; 1000 permutations; cluster-forming threshold α = 0.05)^37^. Effect sizes (Cohen’s d) were computed for all significant contrasts.

### 2.7. Classification Analysis

To assess the predictability of SO occurrence and subtype before onset, we extracted a comprehensive feature set from the 2-second window preceding the SO trough. Features included normalized band-power measures, inter-band power ratios, and cross-frequency coupling metrics (PAC and AAC). Feature discriminability was first quantified using the area under the curve (AUC) from receiver operating characteristic (ROC) analyses^38^. This ranking process identified theta and delta power as the strongest predictors of SO activity and propagation. Subsequently, logistic regression classifiers were trained using these optimal power features to perform two discrimination tasks: (1) SO detection (real SOs vs. surrogate events) and (2) Subtype differentiation (Global vs. Frontal vs. surrogate events). We deliberately selected a simple, linear classifier to prioritize feature interpretability and to demonstrate the robustness of the theta biomarker, rather than to engineer a complex model for maximal classification performance. We employed a leave-one-subject-out cross-validation scheme with a hybrid calibration step; for each fold, the classifier was trained on the pool of N-1 subjects plus a calibration subset (the first 100 events after sleep onset) from the test subject, and then evaluated on the remaining unseen events. Classifier performance was evaluated using accuracy, sensitivity (true positive rate), specificity (true negative rate), precision (positive predictive value), and F1-score (harmonic mean of precision and sensitivity)^39^.

## 3. Results

The data below progress from the physiological validation of SO subtypes to the characterization of their precursors and, finally, to their predictive utility. We begin by confirming that our detection and clustering framework reproduces canonical SO properties, such as a decreasing trend in density and in the magnitude of the negative peak amplitude along the anteroposterior axis, as well as coherent propagation patterns for Global events. We then systematically examine the two-second pre-onset window, starting with instantaneous spectral power to identify dominant frequency-specific precursors, followed by cross-frequency coupling analyses to characterize the distinct network states that facilitate widespread versus spatially restricted propagation. Finally, we integrate these identified features into a classification model to quantify their ability to predict both SO emergence and spatial extent.

### 3.1. Slow Oscillations Exhibit Canonical Topographies and Distinct Spatial Subtypes

We first confirmed that our detection pipeline aligned with prior research and reproduced the known spatial properties of human SOs. Figure 1A shows an example SO waveform from the raw EEG at channel Fz, demonstrating clear correspondence with the traditional definition of an SO^10^. Across subjects, SO density exhibited the expected anteroposterior gradient during NREM sleep (Figure 1B)^8,10,29^, with the highest rates at prefrontal and frontal electrodes and progressively lower rates over central, parietal, and occipital regions. SO negative-peak amplitudes displayed an analogous pattern^29^, with the largest deflections in frontal areas and diminishing amplitudes posteriorly (Figure 1C). These gradients were present in both N2 and N3 sleep, as shown by SO density (Supplementary Figures S1A–B) and SO amplitude (Supplementary Figures S1C–D). Together, these spatial profiles confirm that TP9/TP10 referencing preserved physiologically meaningful gradients despite the absence of mastoid channels.

**Figure 1.**
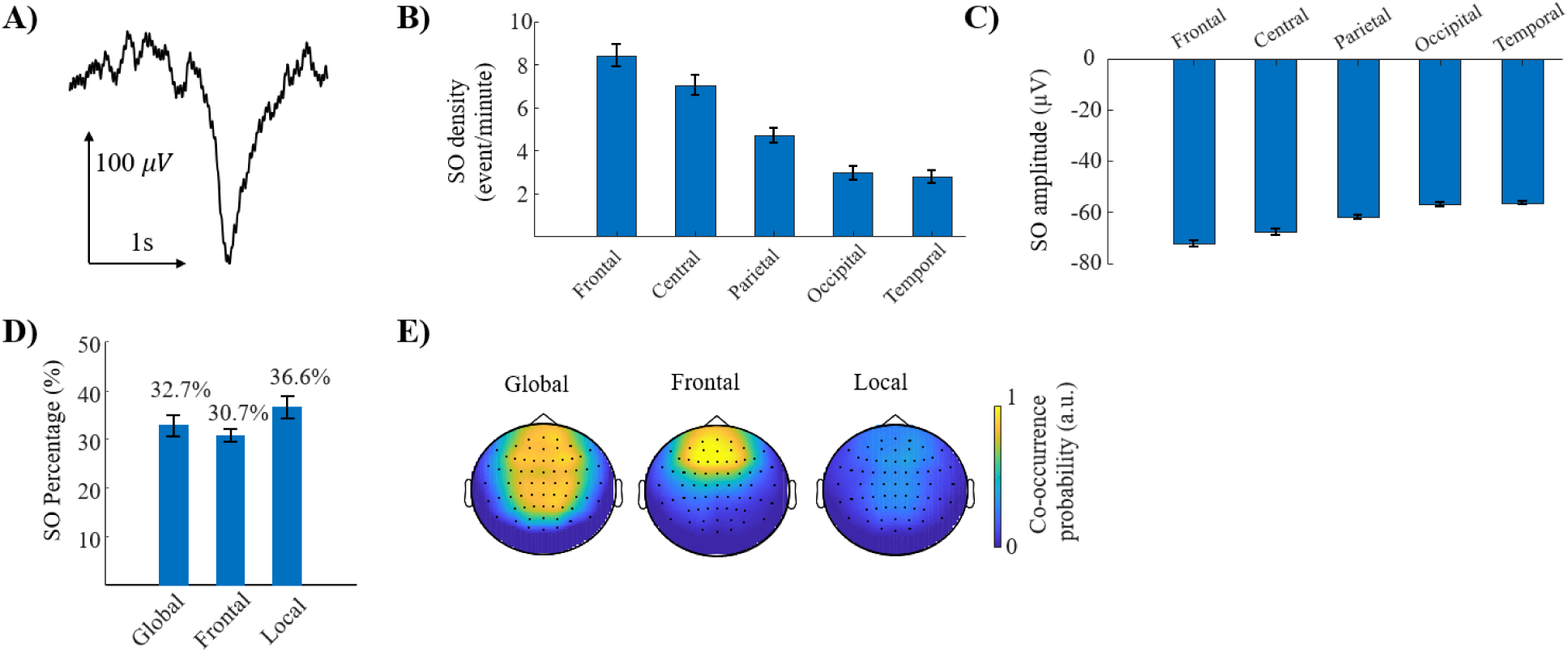
Validation of SO detection and subtype categorization. **A)** Example SO detected in stage N3 (channel Fz, subject 1), showing a characteristic large negative deflection followed by a positive rebound. **B)** Average SO density (events/minute) across scalp regions during NREM sleep. Density declines systematically from frontal to posterior regions, reflecting known SO generation gradients. **C)** Average SO negative-peak amplitude during NREM sleep. The magnitude decreases from frontal to posterior regions, confirming realistic spatial attenuation of SO strength. **D)** Distribution of SO subtypes (Global, Frontal, Local) across all subjects. **E)** Topographic maps of SO subtype co-occurrence, showing the probability that each SO subtype engages a given electrode within a 400ms window. Global SOs exhibit broad, bilateral propagation; Frontal SOs remain confined to prefrontal/frontal channels; Local SOs lack coherent spatial continuity.

Applying k-means clustering to SO co-occurrence patterns separated detected events into three robust subtypes—Global, Frontal, and Local. Figure 1D shows the proportion of SOs assigned to each subtype, and Figure 1E presents topographical maps illustrating the probability of channel involvement in each class. Global SOs engaged widespread bilateral regions spanning frontal, central, parietal, and to a lesser extent occipital cortex; Frontal SOs were confined to prefrontal and frontal electrodes; and Local SOs appeared sparsely without coherent spatial spread. This subtype structure was stable across subjects and provided a principled framework for testing whether distinct pre-onset neural states predict the spatial extent of forthcoming SOs.

Collectively, these results demonstrate that the detection pipeline reproduces canonical SO physiology and supports robust, physiologically meaningful subtype separation.

### 3.2. Theta Power Exhibits a Strong, Early, and SO-Specific Pre-Onset Increase

We next examined the pre-onset dynamics of instantaneous power of frequency bands aligned to the SO trough. Figure 2A illustrates the analysis window relative to an example SO and outlines the workflow for extracting instantaneous power from bandpass-filtered signals. Both Global and Frontal SOs showed a clear elevation in theta power beginning nearly two seconds before the trough, and this increase was substantially stronger than that observed in surrogate events (Figure 2B). Statistical comparisons with surrogate baselines confirmed that theta power rose significantly prior to both SO types (Global: t(28)=7.76; Frontal: t(28)=6.59; both FDR < 0.001), establishing theta enhancement as a robust physiological precursor of SOs rather than a reflection of nonspecific EEG fluctuations.

**Figure 2.**
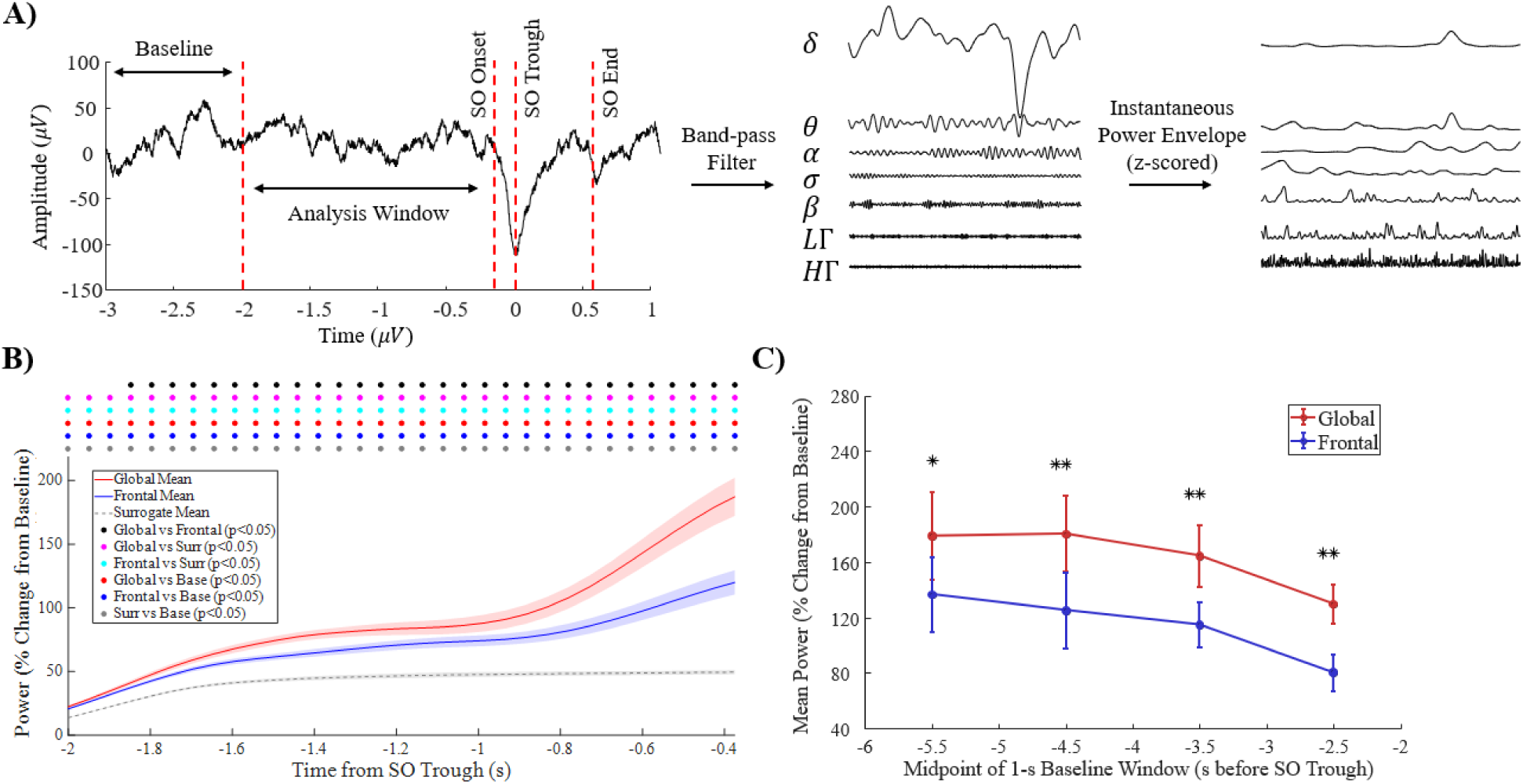
Prediction of SO emergence and spatial extent from pre-onset theta dynamics. **A)** Schematic of the analysis pipeline for pre-onset power. This panel shows the raw EEG signal (including the same SO illustrated in Figure 1A) from 3 s before to 1 s after the SO trough, along with band-pass–filtered waveforms across the frequency bands used in this study. Instantaneous power envelopes derived from these filtered signals are also shown, z-scored to facilitate comparison across bands. Vertical dashed red lines mark the baseline window (2-3 s before the trough), the analysis window (2 s before the trough to the SO onset; on average = −0.37 s before the trough), the SO onset, the trough, and the end of the SO cycle. **B)** Group-averaged theta power time courses (percent change from baseline) aligned to the SO trough for Global SOs, Frontal SOs, and surrogate events. Both SO subtypes show a clear pre-onset increase in theta power compared to surrogate activity, but Global SOs exhibit a stronger and more sustained rise (average t = 5.01, FDR < 0.05), indicating that enhanced pre-onset theta selectively characterizes widespread propagating SOs. Colored dots positioned above the traces denote time bins with significant differences for pairwise comparisons: black (Global vs. Frontal), magenta (Global vs. Surrogate), cyan (Frontal vs. Surrogate), red (Global vs. Baseline), blue (Frontal vs. Baseline), and gray (Surrogate vs. Baseline). Shaded regions represent SEM. **C)** Sensitivity analysis of theta power across baseline selections. Each point represents the midpoint of a 1-s baseline window used to normalize theta power within the analysis window (panel A; −2 to −0.37 s before SO trough). Asterisks (* p < 0.01; ** p < 0.001) indicate significant differences in mean theta power between Global and Frontal SOs for the corresponding baseline. Moderate-to-strong effect sizes were observed for baseline windows between −6 and −2 s relative to the SO trough, demonstrating that elevated pre-onset theta is a stable and reliable signature of SOs and their subtypes.

Critically, Global SOs exhibited a substantially stronger and more sustained theta increase than Frontal SOs. The Global–Frontal difference was significant from approximately −1.85 to −0.37 s before the trough (FDR-corrected; average t=5.01, peak t=7.60). This anticipatory theta separation remained statistically robust when alternative 1-second baseline windows within the −6 to −2 s interval were used, yielding moderate-to-strong effect sizes (Cohen’s d = 0.45–0.77) across all baselines (Figure 2C); as expected, effect sizes uniformly decreased for baselines farther from the analysis window. Together, these results demonstrate that enhanced pre-onset theta activity is both SO-specific and subtype-specific, providing a stable physiological marker that differentiates whether an upcoming SO will propagate globally or remain spatially restricted.

Of note, although theta power emerged as the dominant discriminator, several additional frequency bands displayed meaningful, albeit less reliable, pre-onset differences (Supplementary Figure S2). Delta power increased prior to both SO subtypes and showed a weak Global > Frontal trend (t(28) = 2.21) relative to baseline. While SOs can contribute to power in the delta-band activity, this increase reflects delta-range fluctuations occurring well before the SO trough and hence distinct from the SO waveform itself. Alpha and beta bands exhibited moderate Global > Frontal separation (t(28) = 3.1 and t(28) = 3.4, respectively), but showed no consistent deviations from baseline. None of these bands approached the temporal specificity, effect magnitude, or subtype separation observed with theta. Sigma, low gamma, and high gamma showed minimal differences and did not reliably distinguish Global from Frontal SOs or from surrogate events. Together, these findings indicate that although several frequency bands display preparatory adjustments, theta is the only frequency band that consistently and robustly predicts both SO emergence and spatial extent.

### 3.3. Cross-Frequency Coupling Reveals Distinct Network States Preceding Global and Frontal SOs

PAC analyses revealed strong differences in the cross-frequency structure of brain activity preceding different SO subtypes. Figure 3A illustrates the intuitive relationship between delta phase and theta amplitude: during the delta cycle, theta amplitude peaks around the down-to-up and up-to-down transitions (approximately 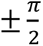), producing a notable MI of 0.0031 for epochs preceding Global SOs. In contrast, epochs preceding Frontal SOs showed a much weaker delta–theta coupling (MI=0.0002). Figure 3B extends this analysis across all frequency-band pairs, showing the t-values for Global–Frontal differences in MI. Consistent with Figure 3A, Global SOs exhibited significantly stronger coupling between delta phase and theta amplitude (t(28) = 2.96; FDR = 0.020). In contrast, Frontal SOs were characterized by significantly stronger coupling between low-frequency phases (theta, alpha) and higher-frequency amplitudes (beta, low gamma). For example, theta–beta coupling was markedly stronger in Frontal than Global SOs (t(28) = −11.98; FDR < 0.001). Baseline comparisons confirmed that this elevated theta- and alpha-phase modulation of high frequencies is a specific feature of the Frontal SO pre-onset state, whereas Global SOs primarily exhibit delta-driven modulation (Supplementary Figure S3). These coupling patterns suggest that Frontal SOs arise from a locally synchronized state dominated by activity at higher frequencies.

**Figure 3.**
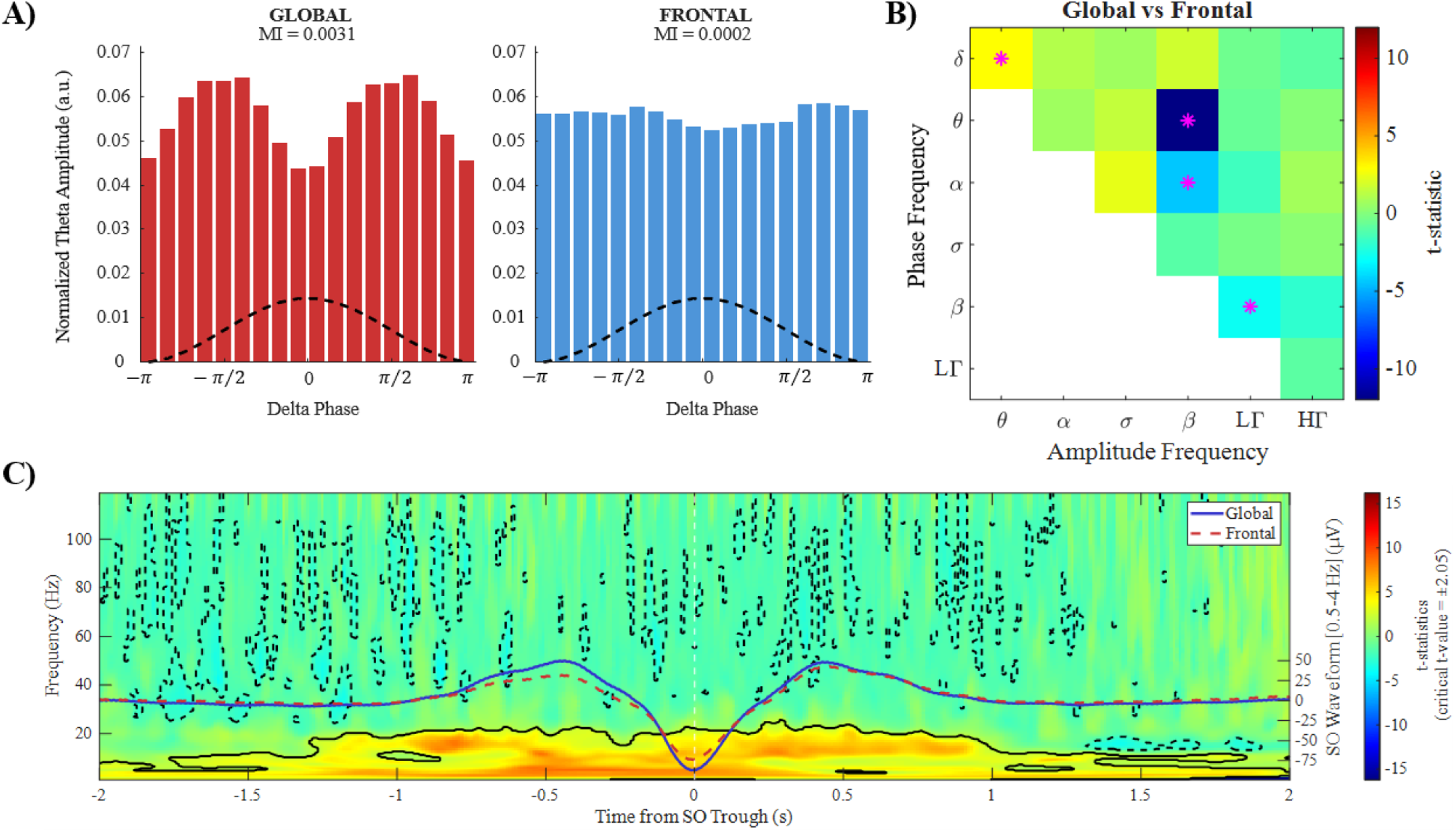
Differentiation of Global from Frontal SOs via cross-frequency and time–frequency dynamics. **A)** PAC analysis using the MI. Global SOs show markedly stronger delta-phase/theta-amplitude coupling than Frontal SOs (t(28) = 2.96; FDR = 0.020). The MI peak for Global SOs occurs at the down-to-up and up-to-down transitions of the delta cycle, indicating coordinated recruitment of theta during global propagation. **B)** PAC matrix comparing coupling between slower-frequency phases and faster-frequency amplitudes. The color bar represents t-values, with significant differences exceeding |t| = 2.05 (FDR-corrected). Global SOs exhibit significantly stronger delta–theta coupling relative to Frontal SOs, whereas Frontal SOs show significantly stronger theta–beta, alpha–beta, and beta–low gamma coupling. These patterns suggest that Global SOs reflect a low-frequency–coordinated state that facilitates long-range propagation, whereas Frontal SOs arise from a locally synchronized, high-frequency–engaged cortical state that may impede large-scale travel. **C)** TFD illustrating differences in spectral dynamics between Global and Frontal SOs. The left y-axis indicates frequency from 1–119 Hz, while the right secondary y-axis indicates the amplitude (*μV*) of the grand-average SO waveforms (0.5–4 Hz) superimposed on the TFD (Global: solid blue line; Frontal: dashed red line), ranging from -75 to 50 *μV*. The x-axis shows time from −2 to 2 s relative to the trough. Solid contours denote significant power increases, and dashed contours denote significant power reductions. Global SOs show broad pre-onset increases across the delta (mean t = 5.17), theta (t = 4.80), alpha (t = 4.17), and sigma (t = 4.45) bands, with 62–98% of cluster pixels surviving permutation-based correction (cluster-level p < 0.05). No enhancement is observed in low- or high-gamma frequencies.

Time–frequency decomposition (TFD) further reinforced the distinction of neural activity in epochs preceding Global vs Frontal SOs (Figure 3C). Compared to Frontal SOs, Global SOs showed broad pre-onset power increases across delta, theta, alpha, and sigma bands (mean significant t-values 4.17–5.17, with 62–98% of cluster pixels surviving permutation-based correction), whereas Frontal SOs exhibited comparatively weaker increases. Supplementary Figure S4 displays the baseline-referenced TFDs: although both subtypes demonstrate significant low-frequency enhancements, Global SOs clearly display broader and stronger involvement in both time and frequency. Additionally, Figure 3C reveals a notable divergence in high-frequency dynamics: Global SOs exhibit a significantly stronger reduction in high-frequency power prior to the SO trough compared to Frontal SOs. Together, these findings, in line with cross-frequency analysis, indicate that Global SOs emerge from a state of enhanced low-frequency coordination and reduced high-frequency activity, whereas Frontal SOs may arise within a more locally synchronized, high-frequency–coupled environment that remains spatially restricted.

AAC analyses revealed that both Global and Frontal SOs involve substantial reconfiguration of cross-frequency amplitude relationships relative to baseline (Supplementary Figure S5). Prior to both SO subtypes, AAC was broadly reduced compared with baseline across low- and mid-frequency pairs—including delta–theta, delta–alpha, theta–beta, and alpha–beta—indicating a general decoupling of cross-frequency co-fluctuations before SO onset. Direct comparisons between Global and Frontal SOs within the pre-onset analysis window showed that Global SOs exhibited significantly greater reductions in AAC between low-frequency bands (delta, theta) and higher-frequency bands (sigma, beta), as well as reduced coupling involving high gamma. In summary, AAC was reduced prior to both SO subtypes relative to baseline, and Global SOs showed significantly greater reductions than Frontal SOs.

### 3.4. Pre-Onset Neural Dynamics Accurately Predict Both SO Occurrence and Spatial Extent

Finally, we assessed whether pre-onset spectral features could predict upcoming SOs and their subtype. Using logistic regression with leave-one-subject-out cross-validation, theta power alone reliably differentiated true SOs from surrogate events, achieving 95.42% accuracy, 93.42% sensitivity, 96.68% specificity, and an F1-score of 0.94 (Table 1). These results demonstrate that the theta precursor provides a robust, physiologically grounded signal for SO detection prior to the trough. When classifying SO subtypes, theta power alone achieved 83.17% accuracy. Surrogate events were identified with excellent precision (F1 = 0.96), consistent with the distinctiveness of SO-specific theta activity. Sensitivity for Global (64.17%) and Frontal (59.37%) SOs was moderate. Including delta power yielded a modest but meaningful improvement: the combined model (theta + delta), showed overall accuracy of 84.74%, Global SO sensitivity of 74.30%, and Frontal SO sensitivity of 61.83%. These classification results establish that pre-onset spectral features can reliably differentiate SOs from background activity and predict their spatial extent using a simple linear model.

**Table 1.**
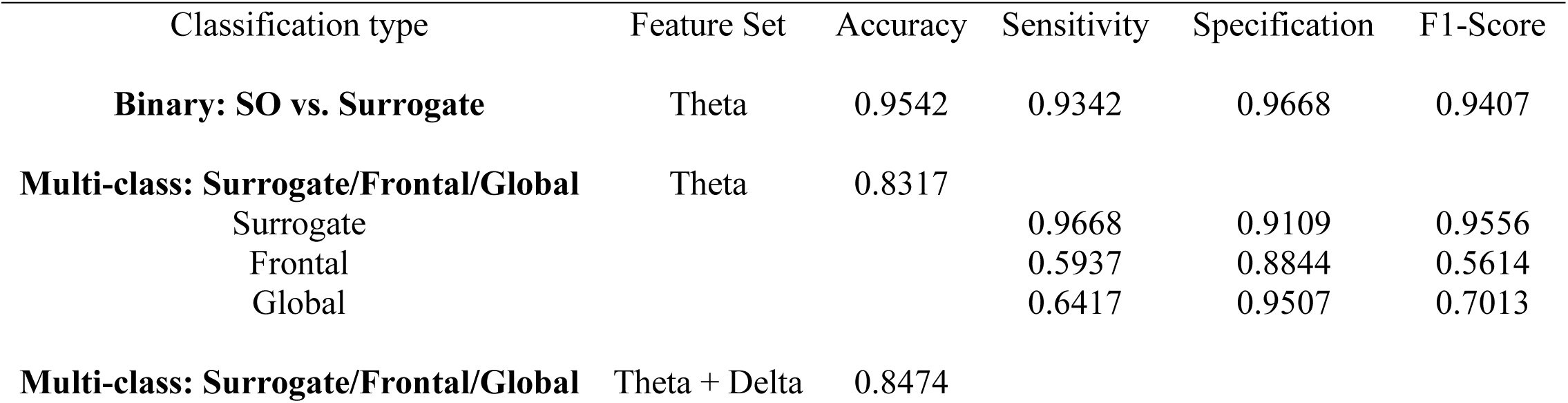
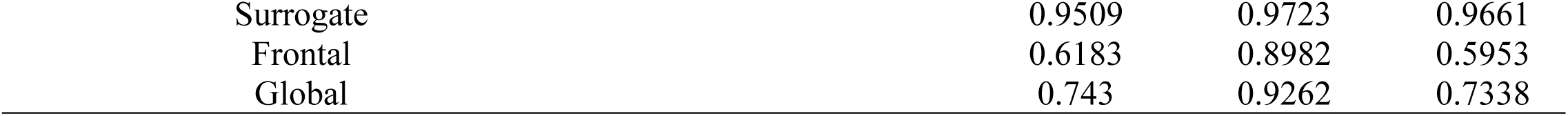
Prediction of SO occurrence and spatial extent using pre-onset spectral features. Predictive performance of linear logistic regression classifiers trained on baseline-normalized spectral features from the 2-s interval preceding SO onset. Theta power emerged as the strongest single predictor across ROC analyses (AUC = 0.93). Overall accuracy, sensitivity, specificity, and F1-score are reported for binary classification (SO vs. surrogate) and multiclass classification (Surrogate, Frontal, Global) using theta power alone or combined theta + delta features. Adding delta power improved multiclass performance, increasing overall accuracy from 83.17% to 84.74% and Global SO sensitivity from 64.17% to 74.30%.

## 4. Discussion

This study demonstrates that SOs during NREM sleep found in the EEG are preceded by structured and measurable neural dynamics. By shifting the analytical focus to the two-second interval before the SO trough, we show that pre-onset spectral features reliably predict both SO occurrence and its spatial extent. Among these features, theta-band activity emerged as a robust and early physiological marker that differentiates SOs from background NREM activity and discriminates Global from Frontal SO subtypes.

A principal finding of this work is the consistent increase in theta power beginning nearly two seconds prior to the SO trough. This increase was present for both Global and Frontal SOs but was significantly stronger and more sustained for Global SOs. Our identification of a theta-band spectral precursor extends prior work by Menicucci et al., who demonstrated that pre-onset sigma-band activity serves as a marker of SO occurrence and amplitude^9^, as well as by Gonzalez et al., who showed that theta-band activity can emerge prior to cortical and thalamic down-states and may contribute to SO initiation^20^. Although we note that Menicucci et al. defined sigma using a broader frequency range (12–18 Hz) than the standard 12–16 Hz band employed here^9,40,41^, our results suggest a functional dissociation: while sigma activity may reflect a generalized thalamocortical entrainment that modulates cortical susceptibility to down-state transitions, theta activity provides the critical information required to predict the spatial extent of the upcoming event. Consistent with this distinction, in our dataset sigma activity exhibited substantially lower predictive capability for SO detection than theta and failed to account for the pronounced pre-onset divergence between Global and Frontal SOs.

The predictive strength of theta also exceeded that of delta power, which showed only modest ability to differentiate Global from Frontal subtypes. This pattern suggests that theta dynamics can capture aspects of network organization preceding an SO event in the EEG that are not fully reflected in slow-wave activity itself. Collectively, these findings indicate that pre-onset theta could reflect a transitional cortical state that precedes large-scale synchronization underlying SO expression. Notably, theta enhancement emerges approximately two seconds before the trough, defining a preparatory window that precedes conventional detection thresholds. This timescale substantially exceeds typical algorithmic and hardware latencies in existing reactive systems, underscoring the practical relevance of theta-based prediction for anticipatory closed-loop neuromodulation.

Importantly, these findings do not imply that pre-SO theta activity is a causal driver of SO initiation. While SOs can recur with some regularity and sometimes appear in clusters, their timing is not strictly periodic, making it less likely that a preceding event’s residual “aftereffects” alone could account for the specific timing and subtype of a subsequent SO. We further reduced concerns about serial dependence by restricting analyses to temporally isolated SOs (excluding events occurring within 1.5 s of a preceding SO trough) and by comparing them against surrogate epochs constructed at least ±3 s from any detected SO trough. Notably, theta power differentiated Global from Frontal subtypes nearly two seconds before the SO trough. This lead time is more consistent with a structured transition between network states linked to subsequent spatial propagation than with a simple consequence of immediate event history. Thus, pre-onset theta appears to reflect a preparatory network configuration for spatial propagation that is at least partly independent of the SO’s immediate temporal context.

The temporal interval identified as significant in the theta-based predictability analysis in this study likely partially overlaps with the early phase of the SO down-state. As such, this window may capture neural activity associated with the initial hyperpolarizing transition rather than exclusively reflecting pre-event background dynamics. This interpretation is consistent with prior work demonstrating that the slope of the transition into the down-state plays a critical role in shaping subsequent up-state coordination and in determining the effective nesting of sleep spindles and hippocampal ripples. Steeper and more spatially synchronized slow waves promote more reliable spindle–ripple coupling, whereas flatter, spatially restricted waves are less effective in driving hierarchical oscillatory organization^42,43^. The present findings therefore suggest that pre-trough theta differences may reflect signatures of distinct hyperpolarization processes that precede SO trough formation and bias the subsequent spatial extent of the oscillation. Future studies should explicitly examine the causal relationship between early theta-band activity and fine-grained SO waveform features—including down-state slope and synchronization efficiency—which are key determinants of sleep-dependent memory consolidation.

While theta power provides the primary predictive signal, time-frequency and cross-frequency analyses revealed that Global and Frontal SOs are preceded by qualitatively distinct pre-onset configurations beyond simple power differences. In the two-second window before the trough, Global SOs exhibited broadband power increases from delta through sigma bands, accompanied by reduced power spanning beta to high-gamma frequencies, decreased AAC among higher-frequency pairs, and stronger delta–theta PAC. Frontal SOs, in contrast, showed more prominent alpha and beta engagement, increased AAC among fast-frequency pairs, and weaker temporal deviations from baseline. These convergent results support the interpretation that widespread propagation is facilitated by a simplified, more decoupled pre-onset network state, whereas Frontal SOs arise from a more interconnected and higher-frequency-engaged environment. The current findings therefore suggest that SO spatial profiles may be linked to preceding neural activity rather than determined solely by post-initiation processes. Importantly, this preceding fingerprint provides crucial information for optimizing SO-targeted closed-loop stimulation approaches.

These converging findings—particularly the robust theta precursor and distinct network configurations preceding different SO subtypes—have direct implications for closed-loop stimulation strategies targeting SOs. Most existing systems rely on reactive detection of SO troughs or down-state transitions, which inherently limits the temporal precision with which stimulation can be aligned to the SO up-state. The robust predictability afforded by pre-onset theta activity suggests that SOs can be anticipated rather than merely detected, potentially enabling stimulation to be delivered more precisely at a desired SO phase without relying only on threshold crossings. Moreover, the ability to differentiate Global and Frontal SOs prior to onset can particularly relevant for selective neuromodulation strategies. Globally propagating SOs have been shown to exhibit stronger coordination with sleep spindles and to be associated with episodic memory, whereas more spatially restricted SOs tend to show weaker spindle associations and no relationship with episodic memory^14,29^. Although optimization of stimulation protocols for targeting slow oscillations is ongoing and various approaches have been proposed^30,44,45^, failure to account for the heterogeneity of SO subtypes may help explain the mixed outcomes observed in studies employing uniform SO modulation strategies, in which physiologically distinct SO subtypes are indiscriminately targeted^46–49^.

In the present study, predictive modeling was deliberately framed in an event-locked manner to establish whether stable and physiologically meaningful pre-onset features exist. Accordingly, our classification analyses assessed whether spectral activity in the two seconds preceding an SO trough contains sufficient information to predict both SO occurrence and spatial extent. From a translational perspective, a complementary and more deployment-oriented approach would involve continuous real-time prediction, in which a sliding temporal window is advanced in small steps across ongoing EEG to estimate the near-term probability of SO occurrence. Such continuous forecasting frameworks are computationally more demanding, but represent a logical next step toward closed-loop implementation. Importantly, the robust and temporally extended theta precursor identified here provides a principled physiological foundation for these approaches, indicating that SO predictability emerges sufficiently early to support real-time prediction schemes in future neuromodulation systems.

Although the present study focused on healthy adults, the results may also inform investigations of pathological sleep dynamics. In disorders such as epilepsy, SOs can become temporally linked with interictal epileptiform discharges during sleep^16,50^. The identification of distinct pre-onset network states raises the possibility that certain SOs may be more permissive to pathological coupling than others, depending on their preparatory dynamics. While the present data do not directly address pathological interactions, the framework established here offers a foundation for future studies examining state-dependent interventions that preserve physiological SOs while avoiding or suppressing pathological ones.

Several limitations should be considered when interpreting our findings. First, the use of TP9/TP10 electrodes as proxy mastoid references likely attenuated absolute amplitude measures; however, the preserved fronto–posterior gradients and canonical SO characteristics support the physiological validity of the detections. Second, the predictive analyses employed linear classifiers to prioritize interpretability and physiological grounding rather than maximal classification performance. Notably, the high accuracy achieved with this simple approach, relying primarily on theta power, demonstrates the robustness of pre-onset theta as a physiological marker without requiring complex feature engineering. Third, although the present analyses employed non-causal filtering and offline classification, the goal of this study was to establish whether predictive neural information exists prior to SO onset rather than to demonstrate a fully deployable real-time system. Importantly, the pre-onset features identified here—particularly sustained theta-band power increases—are compatible with causal filtering and sliding-window implementations^51,52^, suggesting feasibility for future real-time closed-loop applications. Accordingly, while the achieved accuracies demonstrate strong predictability, future work may explore alternative models and causal pipelines better suited for real-time deployment. Finally, the cohort consisted of healthy young adults, and thus the generalizability of these results to older populations or disease states remains to be established.

This study shows that SOs in NREM sleep on the EEG manifold are preceded by structured neural dynamics that can reliably predict both their emergence and spatial propagation. Theta-band power increases nearly two seconds before the SO trough and can serve as a robust marker distinguishing true SOs from background activity, as well as Global from Frontal events. Cross-frequency coupling analyses further demonstrate that widespread SO propagation arises from a distinct preparatory network state characterized by enhanced low-frequency coordination and reduced cross-frequency interference. Together, these findings suggest that the timing and spatial pattern of isolated SO events are tightly related to signatures of network organization preceding SO onset, indicating that the understanding of SO emergence can be enhanced by considering the pre-existing network state. Furthermore, our results provide a foundation for subtype-aware, closed-loop neuromodulation.

