## Supplementary Material for "Predictability of Sleep Slow Oscillation Emergence and Spatial Extent from Pre-Onset Neural Dynamics"

### Tables

**Table S1. Per-participant counts of detected SOs before and after exclusion of SO trains.** Events occurring within 1.5 s of a preceding SO trough on the same channel were excluded to remove SO trains. For each participant, the table reports the total number of detected SOs, subtype-specific SO counts (Global, Frontal, Local), and the number of SOs excluded due to train removal, both overall and separately for each SO subtype.

| Subject ID | Total | Excluded from Total | Frontal | Excluded from Frontal | Global | Excluded from Global | Local | Excluded from Local |
|---|---|---|---|---|---|---|---|---|
| EPCTL01 | 146606 | 31567 | 50364 | 11836 | 35688 | 8644 | 60554 | 11087 |
| EPCTL02 | 200756 | 76859 | 54545 | 20523 | 108593 | 46426 | 37618 | 9910 |
| EPCTL03 | 155883 | 44785 | 59930 | 19125 | 36819 | 11329 | 59134 | 14331 |
| EPCTL04 | 46363 | 9156 | 10354 | 2004 | 11755 | 2611 | 24254 | 4541 |
| EPCTL05 | 135767 | 33161 | 39527 | 10064 | 46636 | 12309 | 49604 | 10788 |
| EPCTL06 | 94475 | 22657 | 39790 | 11074 | 14362 | 3521 | 40323 | 8062 |
| EPCTL07 | 144147 | 44363 | 47878 | 17159 | 64601 | 19059 | 31668 | 8145 |
| EPCTL08 | 118744 | 30694 | 37769 | 10375 | 32516 | 9504 | 48459 | 10815 |
| EPCTL09 | 70967 | 12523 | 17711 | 3582 | 17552 | 4201 | 35704 | 4740 |
| EPCTL10 | 124544 | 35416 | 37736 | 10852 | 43690 | 13724 | 43118 | 10840 |
| EPCTL11 | 205683 | 75401 | 60126 | 22031 | 89786 | 36793 | 55771 | 16577 |
| EPCTL12 | 143534 | 46447 | 38416 | 13118 | 64006 | 23555 | 41112 | 9774 |
| EPCTL13 | 134844 | 37044 | 30198 | 8931 | 51774 | 15525 | 52872 | 12588 |
| EPCTL14 | 110216 | 23976 | 31352 | 8280 | 32788 | 7428 | 46076 | 8268 |
| EPCTL15 | 128694 | 51643 | 46198 | 19835 | 56511 | 24822 | 25985 | 6986 |
| EPCTL16 | 63159 | 11978 | 16367 | 3590 | 9105 | 2240 | 37687 | 6148 |
| EPCTL17 | 125843 | 26020 | 47860 | 11909 | 22760 | 5509 | 55223 | 8602 |
| EPCTL18 | 52169 | 13085 | 13971 | 3627 | 19722 | 5672 | 18476 | 3786 |
| EPCTL19 | 98488 | 25049 | 22208 | 6035 | 45642 | 12222 | 30638 | 6792 |
| EPCTL20 | 53757 | 6782 | 15279 | 2157 | 5741 | 835 | 32737 | 3790 |
| EPCTL21 | 180154 | 52949 | 48383 | 15747 | 69003 | 21697 | 62768 | 15505 |
| EPCTL22 | 51906 | 7883 | 12913 | 2079 | 12152 | 2232 | 26841 | 3572 |
| EPCTL23 | 43214 | 6266 | 8868 | 1376 | 8689 | 1340 | 25657 | 3550 |
| EPCTL24 | 78977 | 15540 | 22887 | 4723 | 22876 | 5381 | 33214 | 5436 |
| EPCTL25 | 108193 | 30350 | 25642 | 8252 | 47580 | 14007 | 34971 | 8091 |
| EPCTL26 | 103649 | 27350 | 36603 | 10430 | 26915 | 7975 | 40131 | 8945 |
| EPCTL27 | 136672 | 45263 | 38544 | 13451 | 49863 | 19107 | 48265 | 12705 |
| EPCTL28 | 80878 | 16333 | 17605 | 3734 | 32992 | 7251 | 30281 | 5348 |
| EPCTL29 | 124842 | 28351 | 38326 | 9458 | 31682 | 7562 | 54834 | 11331 |

### Figures

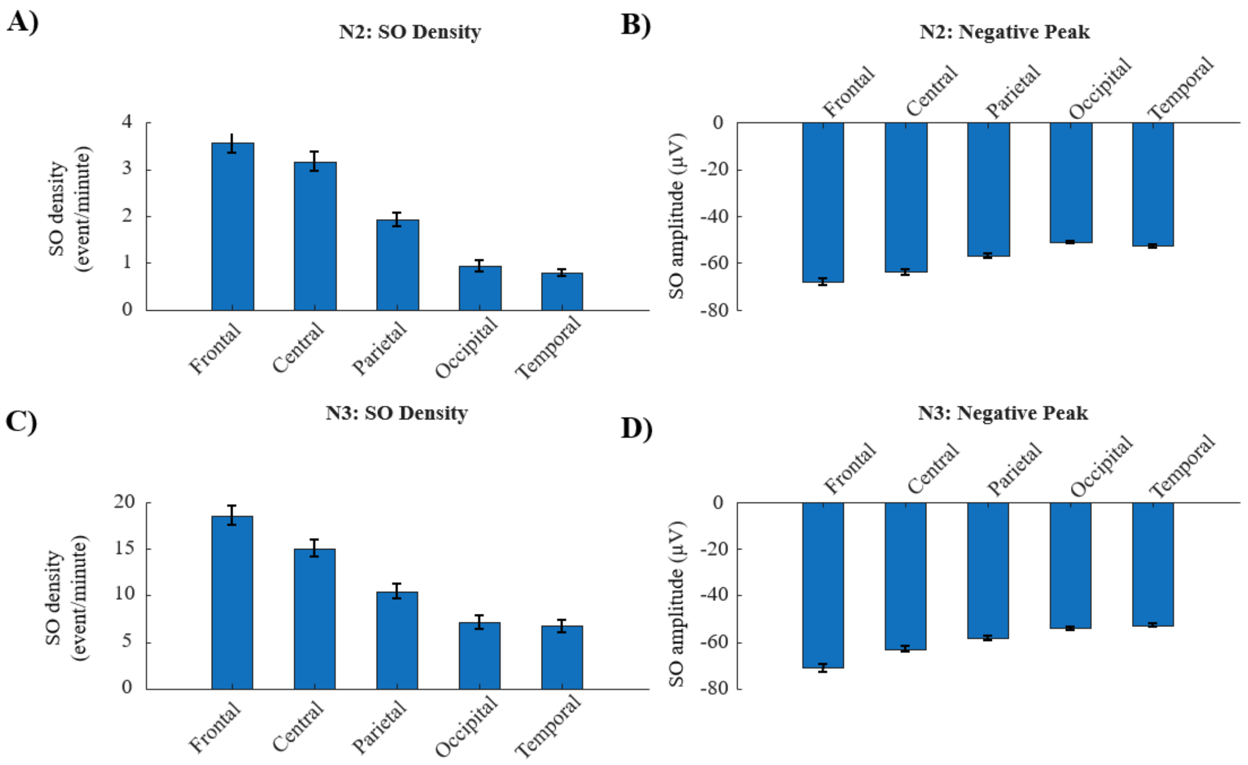


**Figure S1. SO density and amplitude across scalp regions in stages N2 and N3. A–B)** N2: SO density (events/min) and negative-peak amplitude show strong fronto–occipital gradients. **C–D)** N3: The same gradients remain but with larger overall SO densities and amplitudes, consistent with deeper sleep physiology. These patterns confirm that the detection pipeline faithfully reproduces canonical SO spatial organization across NREM stages.

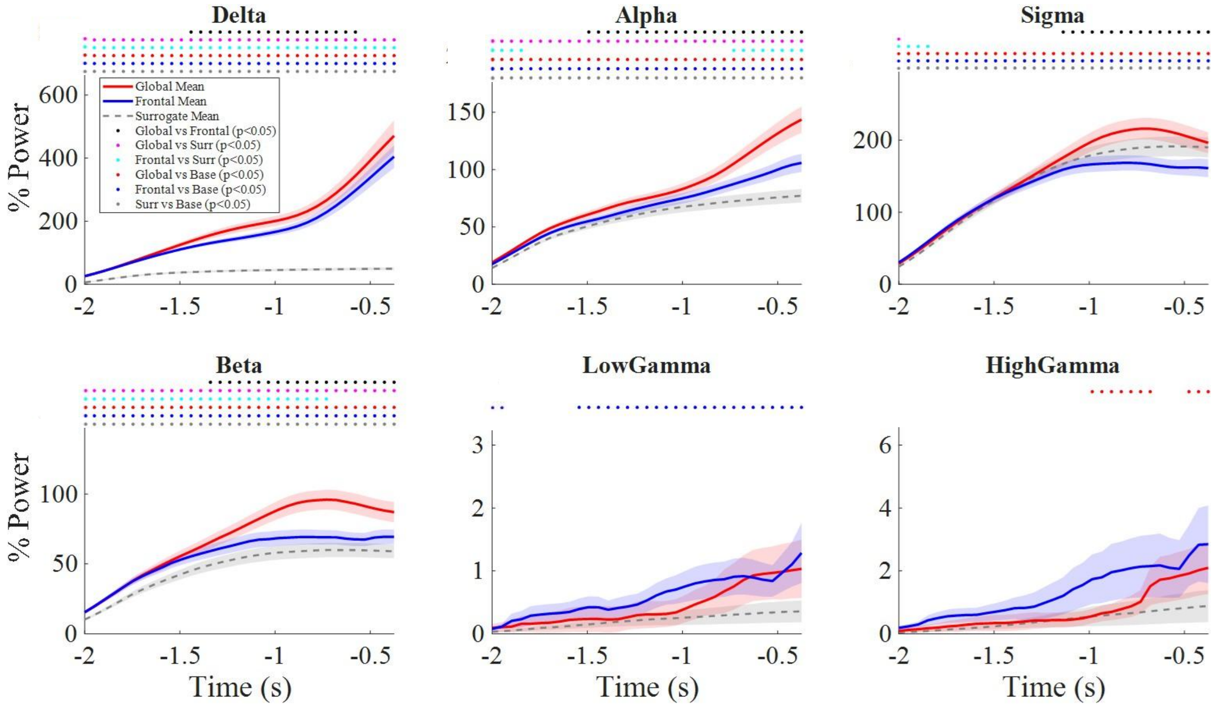


**Figure S2. Frequency-band power dynamics prior to SO onset.** Time courses of delta, alpha, sigma, beta, low gamma, and high gamma power during the −2 to 0 s pre-onset interval for surrogate, Frontal, and Global SOs. None of these bands exhibit the robust, subtype-specific separation seen in theta. Delta shows only weak Global > Frontal differences (t-value = 2.21), and higher-frequency bands fail to distinguish subtypes. Results highlight theta as the only frequency band with reliable and graded anticipatory differentiation.

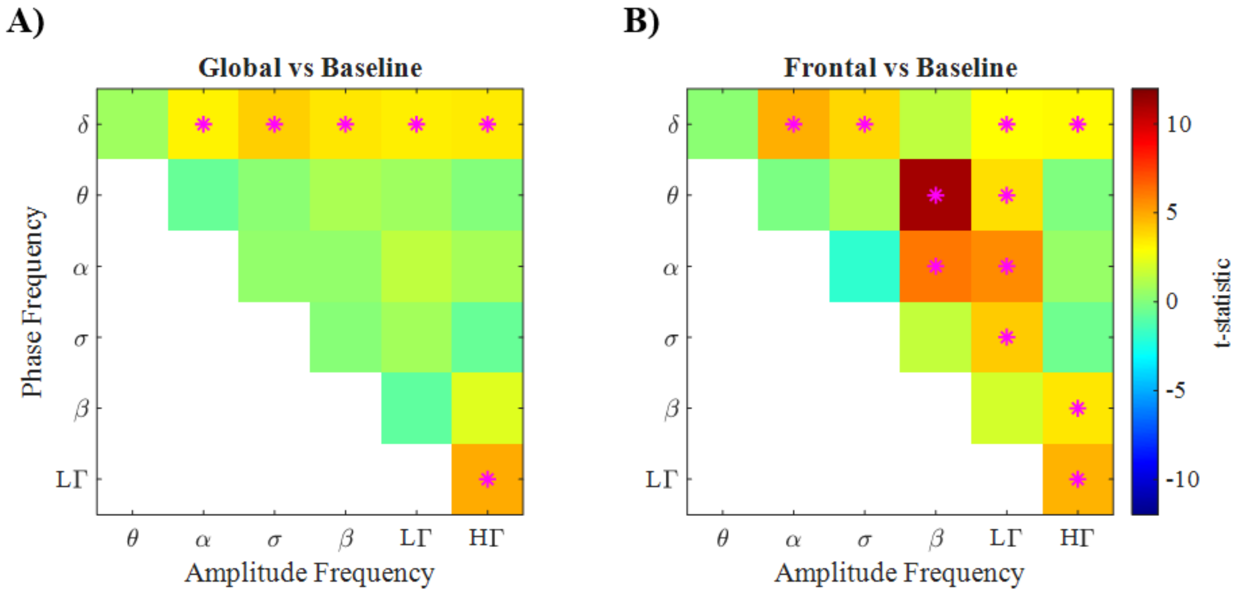


**Figure S3. PAC relative to baseline for Global and Frontal SOs.** PAC matrices comparing each SO subtype to its baseline (−3 to −2 s). Both Global and Frontal SOs show significant increases in delta-phase coupling with alpha, sigma, beta, and gamma amplitudes. Delta–theta PAC remains unchanged from baseline for both subtypes. Frontal SOs additionally show pronounced increases in theta- and alpha-phase coupling to beta and low-gamma amplitudes. These patterns suggest that Frontal SOs arise from a more high-frequency–engaged pre-onset network state, whereas Global SOs do not exhibit elevated high-frequency modulation prior to onset.

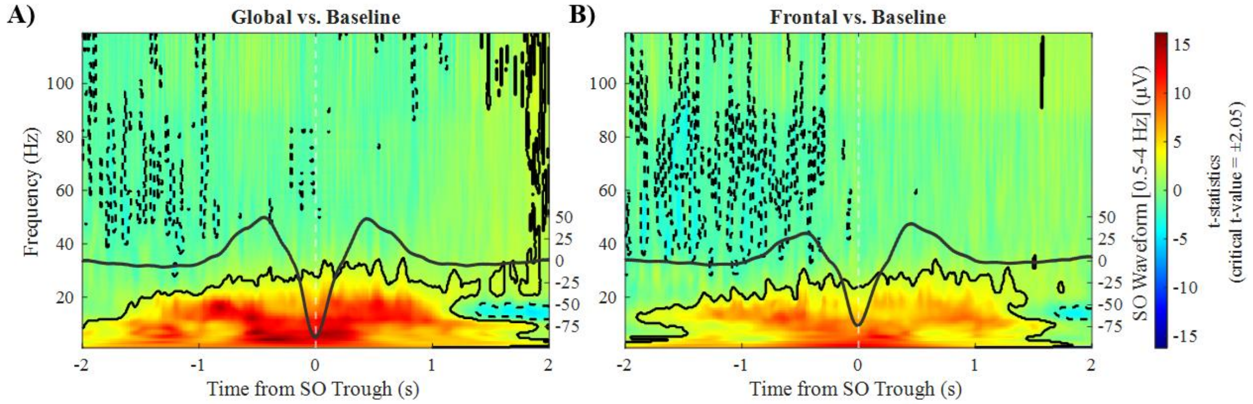


**Figure S4. Time–frequency power relative to baseline for Global and Frontal SOs.** Cluster-corrected time–frequency maps for each SO subtype compared to baseline. Both SO types show significant low-frequency increases prior to trough, but Global SOs exhibit a broader and stronger pattern, particularly in delta and theta bands (Global mean t-value: delta = 8.54, theta = 8.65). High-frequency (beta/gamma) ranges show no significant enhancement, indicating that SO generation and propagation predominantly involve low-frequency mechanisms.

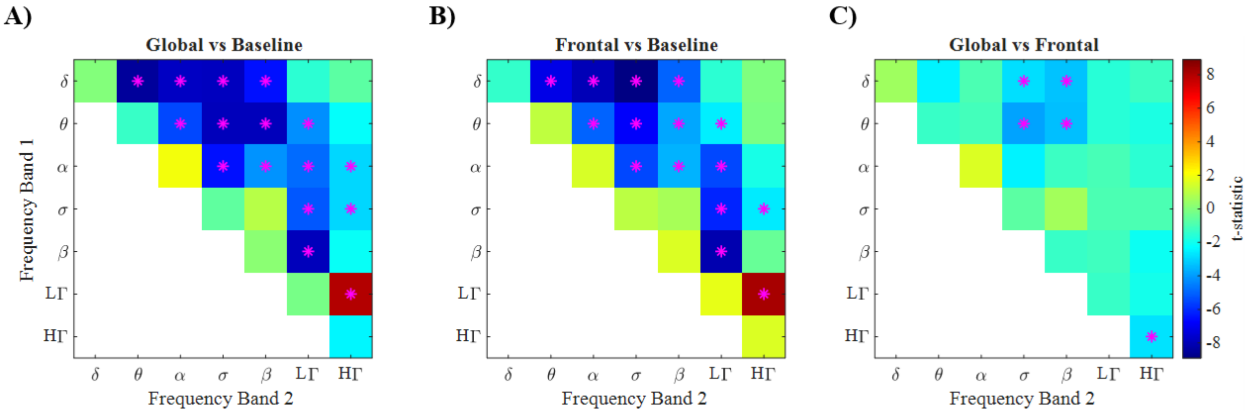


**Figure S5. AAC dynamics across SO subtypes. A–B)** AAC matrices for Global and Frontal SOs relative to the baseline window (−3 to −2 s), computed as the Pearson correlation between Hilbert amplitude envelopes of all frequency-band pairs. Both SO types show a significant reduction in coupling between many low-/mid-frequency combinations (e.g., delta–theta, delta–alpha, theta–alpha) compared to baseline, consistent with a reconfiguration of cross-frequency interactions before SO emergence. **C)** Direct comparison of AAC between Global and Frontal SOs. Global SOs exhibit lower coupling between low-frequency bands (delta, theta) and faster bands (sigma, beta) than Frontal SOs, with differences reaching statistical significance for several pairs (e.g., delta–theta and theta–beta; FDR-corrected $p < 0.05$). These patterns indicate that Global SOs arise from a more spectrally decoupled pre-onset state than Frontal SOs, which may reflect distinct preparatory network configurations associated with broader versus more localized SO propagation.
